# Multi-omics Identify Serotonin Transporter as a Promising Therapeutic Target for Essential Tremor

**DOI:** 10.1101/2024.03.18.585649

**Authors:** Lingbing Wang, Yanjing Li, Suzhen Lin, Zhuofan Zhou, Shaoyi Zhang, Tian-Le Xu, Xing-Lei Song, Yiwen Wu

## Abstract

Essential tremor (ET) stands as one of the most prevalent cerebellar movement disorders. However, effective treatment remains elusive, largely due to a limited understanding of its molecular pathology. Harmaline-induced tremor in mouse is a well-established animal model for ET, while with enigmatic mechanism. The aim of this study was to get insight into the molecular intricacies underlying cerebellar dysfunction in harmaline-induced tremor. Combining LC-MS/MS and RNA-Seq analysis, we delved into the variation of the cerebellum between harmaline-induced tremor and the control ones. This comprehensive investigation revealed a profile of this mouse model from mRNA and protein level, highlighting 5194 correlated coding molecules, with 19 proving to be significant. Further KEGG enrichment analysis identified cerebellar serotonin transporter (SERT) as the key molecule in harmaline-induced tremor. The implications of this transcriptomic and proteomic exploration underscore the potential therapeutic value of targeting SERT as a novel treatment approach for ET. In general, our study unveils crucial insights that could pave the way for molecular target identification and effective therapeutic interventions for ET.

## Introduction

Essential tremor (ET) is one of the most common movement disorders^1,2^, characterized by the rhythmic oscillation of agonist and antagonist muscle groups, typically occurring at a frequency of 8 to 12 Hz^3,4^. The incidence of ET has been reported to be around 0.9%, with a significant increase in prevalence among individuals over 65 years old, reaching 4.6%^5^. Drugs with established efficacy include propranolol, a β-adrenergic blocker, and primidone, an anticonvulsant. However, these medications are associated with side effects, and their efficacy is limited, resulting in an average tremor reduction of only around 50%. Consequently, there is pressing need for targeted therapies that explore new biological pathways. Nevertheless, the underlying mechanisms of ET remain elusive. Harmaline-induced tremor is a recognized model used to screen new therapies for ET^6^. In laboratory animals, a tremor at a frequency of 8-12 Hz can be generated after intraperitoneal injection of harmaline at doses ranging from 10 to 50 mg/kg, with a time latency of 3-10 minutes^7^. Furthermore, Louis *et al*., discovered elevated concentrations of harmane, another compound of harmala alkaloids, in ET patients compared to controls in the Faroe island^8^. This suggests that β-carboline alkaloid may contribute to ET development in both patients and laboratory animals. However, the exact pathology underlying harmaline-induced tremor has not been conclusively elucidated^9^.

The prevailing viewpoint suggests that harmaline can instigate abnormal activation of climbing fibers (CF) within the inferior olivary nucleus (ION), leading to the formation of aberrant synapses with Purkinje cells (PC) in the cerebellar cortex^10^. While recent findings indicate that harmaline induces a burst pattern of activity in Purkinje cells, and the absence of Purkinje cell neurotransmission can attenuate harmaline-induced tremor^11^. This underscores the crucial role of the cerebellar cortex itself, particularly the Purkinje cells, in harmaline-induced tremor. Understanding the mechanism underlying harmaline-induced tremor in the cerebellum may help reveal the molecular basis of ET.

Transcriptomic analysis has been widely recognized as an efficient method for unveiling tissue-specific alterations, including gene splicing, structural variations and transcriptional changes^12^. This technique is capable of uncovering variations in gene expression^13^. However, the significance of post-transcriptional modifications and protein turnover in determining protein function should not be underestimated^14^. The unique advantage of combined analyses of transcriptomics and proteomics lies in their integration, which helps mitigate systematic errors associated with each method individually. In this study, we employed a comprehensive approach integrating both transcriptomic and proteomic analysis to thoroughly disclose the alteration in the cerebellum induced by harmaline treatment. To gain deeper insights into the mechanism contributing to harmaline-induced tremor in the cerebellum, we conducted a comparative analysis of the cerebellums of mice treated with harmaline and their control counterpart. Through the integration of transcriptomic results and proteomic traits, numerous candidate genes were identified. Subsequent extended biological experiments were conducted to validate the role of the candidate molecule serotonin transporter (SERT) in cerebellar PC activity during harmaline-induced tremor. Thus, this study revealed a novel molecular mechanism for harmaline-induced ET and screened potential therapeutic targets.

## Materials and methods

### Tremor detection in freely moving mice

Tremors were recorded using a tremor detector (Medusa, bio-signal), by pre-implanting an electrode slice on the skull of the mice. During tremor measurement, the electrode slice was connected to a transformer, enabling the recording of behavioral data and transforming vibratory signals into digital signals for subsequent analysis. The collected data were analyzed using a preset program that employed the power spectrum density function and further transformed it into frequency domains. Normalization of the spectrum data was achieved through logarithmic (lg) calculation. Adult mice were administered 30 mg/kg harmaline (Topscience, T2792) via intraperitoneal injection (i.p.). For specific treatments, mice were administered DSP-1053 (10 mg/kg), harmaline (30 mg/kg) or an equal amount of saline as a control. DSP-1053 was applied to mice (i.p.) 1 hour before harmaline, ensuring that the blood concentration of serotonin would almost reach its peak when treated with harmaline^15^. The mice were then sacrificed 30 min after harmaline injection.

### Animals

The study was conducted in accordance with the approval of the Animal Ethics Committee of the (Shanghai, China). Analyses were conducted on 6-week-old C57BL/6 male mice, which were subjected with harmaline and DSP-1053 injections, along with their control wild-type littermates (WT). The mice were housed under controlled conditions of lighting (12-hour light, 12-hour dark cycle) and temperature (22 ± 2°C), with unrestricted access to food and water. 30 minutes after injection, animals were anesthetized with isoflurane and subsequently euthanized by decapitation. Cerebellar cortexes samples were promptly dissected and either flash-frozen in liquid nitrogen for proteomic analysis or preserved in RNALater (Beyotime, R0118) for transcriptomic sequencing afterwards.

### Experimental design

In the proteomic analyses, three mice were included in each group (harmaline-induced tremor vs WT). Cerebellar cortex samples were collected, and proteins were extracted, reduced and alkylated. Subsequently, each fraction underwent trypsin digestion to generate peptides. LC-MS/MS spectrometry analysis was performed to identify and quantity proteins, and the resulting data were subjected to the final bioinformatics analysis. For transcriptomic analyses, an equal number of 3 mice were utilized in each group. Following RNA extraction and verification, cDNA libraries were constructed and sequenced to obtain data for the final bioinformatics analysis.

### Protein extraction and digestion and protein-protein interaction analysis

The protein samples were dissociated by SDT buffer (4% SDS, 100 mM Tris-HCl,1 mM DTT!pH 7.6), and the quantity of which were determined by BCA (Bio-Rad, USA). Filter-aided sample preparation (FASP) procedure was used to digest protein with trypsin^16,17^, then the samples were desalted by C18 Cartridges (Empore™ SPE Cartridges C18, Sigma), concentrated and reconstituted by formic acid.

The IntAct molecular interaction database (http://www.ebi.ac.uk/intact/) and STRING software (http://string-db.org/) was used to analyze the protein–protein interaction (PPI) information. Cytoscape software (http://www.cytoscape.org/, version 3.2.1) was then utilized for the PPI visualization.

### RNA extraction and transcriptomic analysis

Total RNA was extracted from cerebellum cortex samples preserved in RNALater using TRIzol reagent. The RNA concentration was quantified at 260 nm using the RNA 6000 Nanodrop on the Agilent 4150 Bioanalyzer (Agilent). Afterwards, mRNAs were isolated using beads with Oligo (dT) and subjected to random fragmentation buffer. Complementary DNAs (cDNAs) were then synthesized based on the extracted mRNA and purified by AMPure XP beads. The enriched cDNAs were further amplified via PCR.

### Liquid chromatograph triple quadrupole mass spectrometer analysis

Q Exactive mass spectrometer (Thermo Scientific) was used to analyze Liquid chromatograph triple quadrupole mass spectrometer (LC-MS/MS) coupled with Easy nLC (Proxeon Biosystems, now Thermo Fisher Scientific). Peptides was dissolved in buffer A (0.1% Formic acid) loading into a a reverse phase trap column (Thermo Scientific Acclaim), the column was connected to a C18-reversed phase analytical column (Thermo Scientific Easy Column). Buffer B (84% acetonitrile and 0.1% Formic acid) was used to separate the peptides with the rate of 300 nL/min.

The mass spectrometer was operated under positive ion mode, the most abundant precursor ions were acquired by HCD fragmentation scanning. The dynamic exclusion duration was set at 40.0 s, survey scans were acquired at a resolution of 70,000 at m/z 200, and the resolution for HCD spectra was set at 17,500 at m/z 200, the isolation width was 2 m/z.

### Bioinformatic analysis

#### GO annotation

NCBI BLAST+ client software were utilized to search the differentially expressed proteins sequences, after the homologous sequences were identified with InterProScan, the protein sequences were mapped with gene ontology (GO) terms, annotated by Blast2GO software, and subsequently plotted by R scripts.

#### KEGG annotation

After completing the annotation, Kyoto Encyclopedia of Genes and Genomes (KEGG) orthology was identified by KEGG database (http://geneontology.org/) with a BLAST search, and then mapped to pathways.

### Western blot

Fresh cerebellar cortex samples from mice were solubilized in lysis buffer containing proteinase inhibitors (Thermo Fisher, A32965) and phosphatase inhibitors (Thermo Fisher, 78420), After sonication and centrifugation, the supernatant was combined with loading buffer (Thermo Fisher, AM8547). Protein concentration was determined using the Pierce BCA Protein Assay Kit (Thermo Fisher, 23225). Following sample preparation, the proteins were loaded onto a 10% SDS-PAGE gel and transferred onto a PVDF membrane (Millipore). The membrane was blocked with 3% BSA (BioFroxx, 4240GR100), and then incubated with primary antibodies: SERT (1:1000, Abcam, ab102048) and GAPDH (Thermo Fisher, A300-639A-T). After an overnight incubation with the primary antibodies, the respective secondary antibodies (1:2000) were applied. Signals were detected using Tanon-5200 system (Tanon).

### ELISA assay

An ELISA kit (MM-0443M1) was used to determine the concentration of serotonin in the cerebellum of harmaline-treated mice and their corresponding control counterpart. The cerebellum samples were washed, sonicated, and centrifuged with PBS (pH=7.4, Sangon biotech, B548117-0500), and the resulting liquid supernatant was collected for further examination. The results were measured using a multi-mode microplate reader.

### Primary culture of cerebellar cortical neuron

At embryonic day 18 (E18), mice were anesthetized with diethyl ether. Following sterilization with 75% ethyl alcohol, a midline incision was made in the abdomen to expose and separate the uterus form the pregnant mice. Next, fetal mouse heads were extracted and placed in a Petri plate containing dissection buffer (DMEM + 5% penicillin/ streptomycin). The scalps of the fetal mice were then cut open using sterilized scissors, and the brain tissues were carefully extracted. Under an optical microscope, the cerebellum was separated from the whole brain tissue, the meninges were peeled off from the surface of the brain. The isolated cerebellum was placed in a 7 mL centrifuge tube with digestion buffer (2 mL 0.25% trypsin + 2 mL dissection buffer). The centrifuge tube was transferred to a constant-temperature and humidity incubator for 15 minutes, with gentle shaking every 5 minutes during incubation. After incubation, excess digestion buffer was removed and 4 mL of complete culture medium (DMEM + 10% FBS + 1% GlutaMAX) was added. The mixture was then homogenized with the digested brain tissue, filtered through a filter net, and the resulting cell suspension was centrifuged at 1000 rpm for 5 minutes. After centrifugation, the cell pellet was resuspended in 2 mL of complete culture medium. After cell counting, the cell suspension was added to cell-culture dishes preloaded with 2 mL of culture medium for cerebellar neurons (50% complete culture medium + 50% neural selective medium (Neuralbasal + 2% B27 + 1% GlutaMAX) + 2 uL T3 (20 mg/ml)). Cerebellar neurons were cultured in a constant-temperature and humidity incubator.

### Cell electrophysiology

The electrophysiological characteristics of cultured cerebellar cortical neurons were recorded using voltage clamp techniques in the whole-cell mode of patch clamp at room temperature. Data acquisition and analysis were performed using the patch clamp amplifier system (MultiClamp 700B) and digital analog converter (Digidata 1550B and pClamp10). For the recording, glass electrodes filled with filtered electrode fluid (10 mM NaCl, 5 mM KCl, 1 mM MgCl2, 2 mM CaCl2, 10 mM HEPES, 10 mM Glucose, pH 7.4, 310-320 mOsm/L) were utilized. The resistance of the glass electrode, when immersed in the extracellular fluid (150 mM NaCl, 5 mM KCl, 1 mM MgCl2, 2 mM CaCl2, 210 mM glucose), ranged between 2-5 MΩ. Following baseline normalization, the glass electrode was connected to the cultured cerebellar cortical neuron under negative pressure. Then, the cell membrane was aspirated until it broke, achieving high-pressure sealing and establishing the whole-cell mode of patch clamp.

### Virus injection and optical recording

The mouse was anesthetized with isoflurane. Then, the mouse’s head was restrained in a stereotaxic instrument, and the scalp above the cerebellar region was incised following fur removal. Adequate sterilization was applied, and a hole was drilled in the skull. A Hamilton syringe containing the virus was inserted into the mouse cerebellum with the coordinates (relative to Bregma: AP -6.75 mm, ML 1.8 mm, DV -2.5 mm). A total of 300 nL of the virus, administered at a rate of 0.1 uL per minute, was injected into the cortex of the mouse’s cerebellum. Ten minutes after completing the injection, the syringe was gradually removed at a rate of 0.05 mm per minute. Subsequently, an optogenetic fiber (Thinker Tech Nanjing Biotech Co., Ltd) was implanted above the virus-injection area (relative to Bregma: AP -6.75 mm, ML 1.8 mm, DV: -2 mm). Following these procedures, the scalp was sutured, and the mice received appropriate post-operative care. Upon full recovery and viral transgene expression, the optical signal of serotonin was detected using Signal-channel Fiber Photometry (Thinker Tech Nanjing Biotech Co., Ltd).

### Statistical analysis

Tissue level of serotonin system, tremor detection of mice and electrophysiological parameters were evaluated using GraphPad Prism9 (GraphPad Software, La Jolla, CA). A two-tailed Student’s *t*-test was used for the comparison of two groups, while one-way analysis of variance (ANOVA) was used for comparisons involving more than two groups. The homogeneity of variance was assessed using Brown-Forsythe and Bartlett’s tests. All data were presented as mean ± SEM (standard error of the mean), with ‘n’ representing the sample number (i.e., the number of independent experiments or cell numbers). Significant differences were denoted as \**P* < 0.05!\*\**P* < 0.01! \*\*\**P* < 0.001 and \*\*\*\**P* < 0.0001.

## Results

### Rhythmic activity detection of mice induced by harmaline

We established a harmaline-induced tremor model in mice following established protocols from previous studies^6,18,19^ (Fig. 1A). Upon injection with harmaline (dissolved in DMSO and diluted with saline, *in vivo* 30 mg/kg), mice exhibited pronounced tremors manifesting across the head, trunk, and limbs. Compare to their control littermates injected with the vehicle (DMSO diluted with saline), the tremors presented in harmaline-injected mice exhibited a distinct frequency range of 8-20 Hz, initiating approximately 3 minutes after intraperitoneal injection (Fig. 1B-G). Additionally, mice subjected to harmaline treatment displayed significantly higher intensity in the 8-20 Hz range in contrast with the control group (Fig. 1H). The tremors persisted for approximately 2 hours, reaching their peak at 30 minutes post-injection. Subsequently, we sacrificed the model mice 30 minutes after harmaline administration and separated cerebellar cortex for protein and RNA extraction.

**Fig 1.**
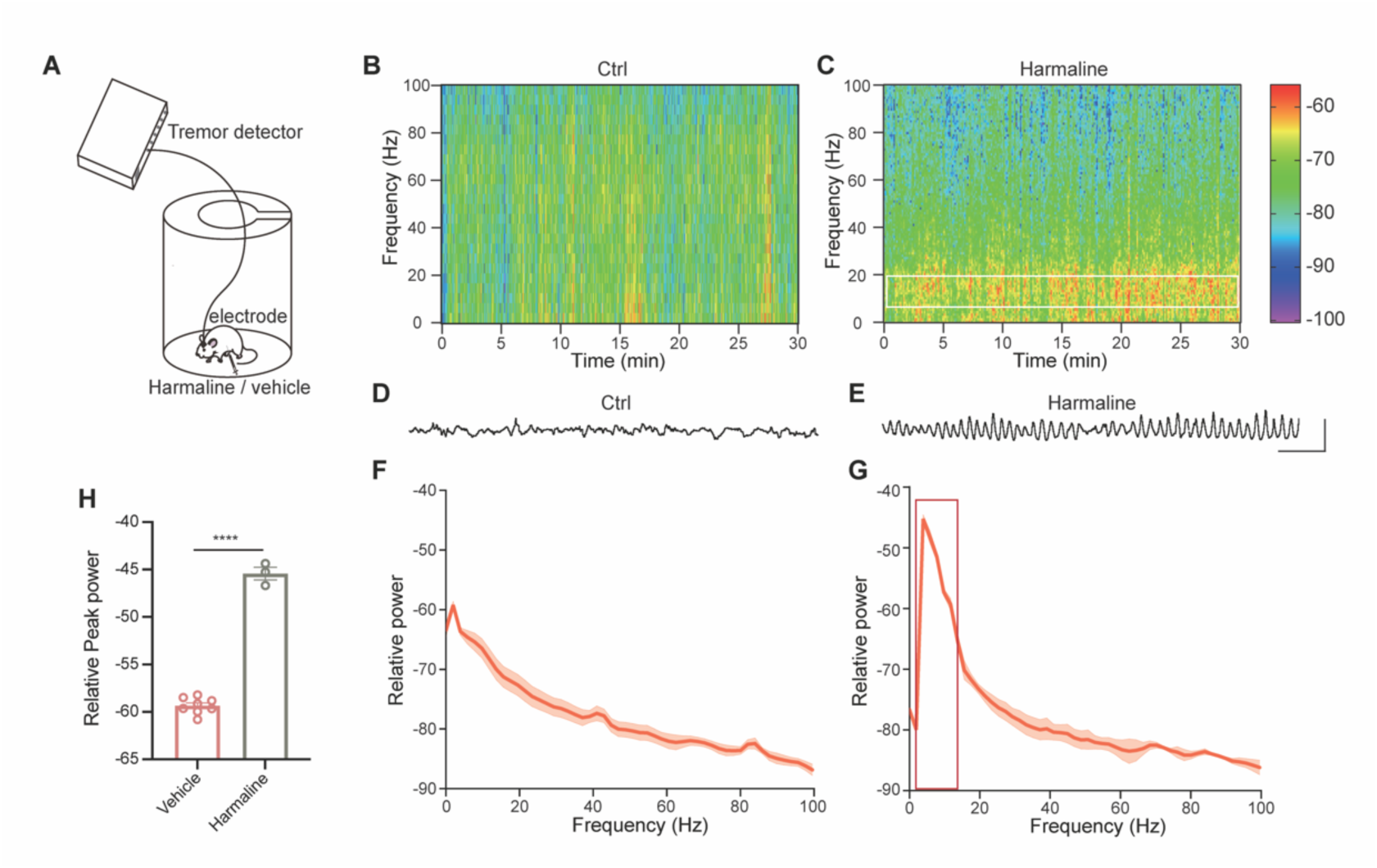
Establishment and phenotypic characterization of the Harmaline-induced mouse tremor model. (**A**) Schematic paradigm for detection of harmaline-induced essential tremor. (**B**-**C**) Representative spectrograph calculated from 30-minute tremor measurements based on head vibration respectively in control group (**B**) and harmaline-exposure group (**C**). (**D**-**E**) Representative raw data of weight measurement using tremor detector based on head vibration in control group (**D**) and harmaline-exposure group (**E**). (**F**-**G**) Averaged power spectrum from data in (**B**) and (**C**). n = 8 for control group (**F**) and n = 3 for harmaline-exposure group (**G**). (**H**) Comparison of averaged peak power from 8 to 20 Hz between control (n = 8) and harmaline-exposure (n = 3) group. Data presented as mean ± SEM, Ctrl, -59.34 ± 0.3005; Harmaline, - 45.43 ± 0.6715. \*\*\*\**P* < 0.0001, two-tailed Students’ *t*-test.

### Transcriptomic and proteomic workflow and overall characterization following harmaline-induced ET model

To date, no study has documented transcriptomic or proteomic changes in cerebellar cortex of harmaline-administrated mice for ET model. As illustrated in the schematic workflow (Fig. 2A), we performed both transcriptomic and a proteomic analysis respectively, comparing 3 mice treated with harmaline to 3 vehicle-treated littermates so as to unravel the transcriptomic and proteomic profile changes associate with harmaline administration. For each cerebellar cortex lysate sample, we performed LS-MS/MS mass spectrometry analysis in triplicate. The reliability of our analysis can be pledged by the high technical reproducibility observed in our experiments and the minimal variation in our cerebellar cortex samples. Notably, for transcriptomic analysis, RNA samples met all the criteria for cDNA library construction standard (all OD260/280=2.1, RINs (RNA Integrity Number) vary from 8.9 to 9.2). As for proteomic analysis, all electrophoretic bands were clear and the volumes of each sample were sufficient (1431.6 μg ∼ 2555.6 μg).

**Fig 2.**
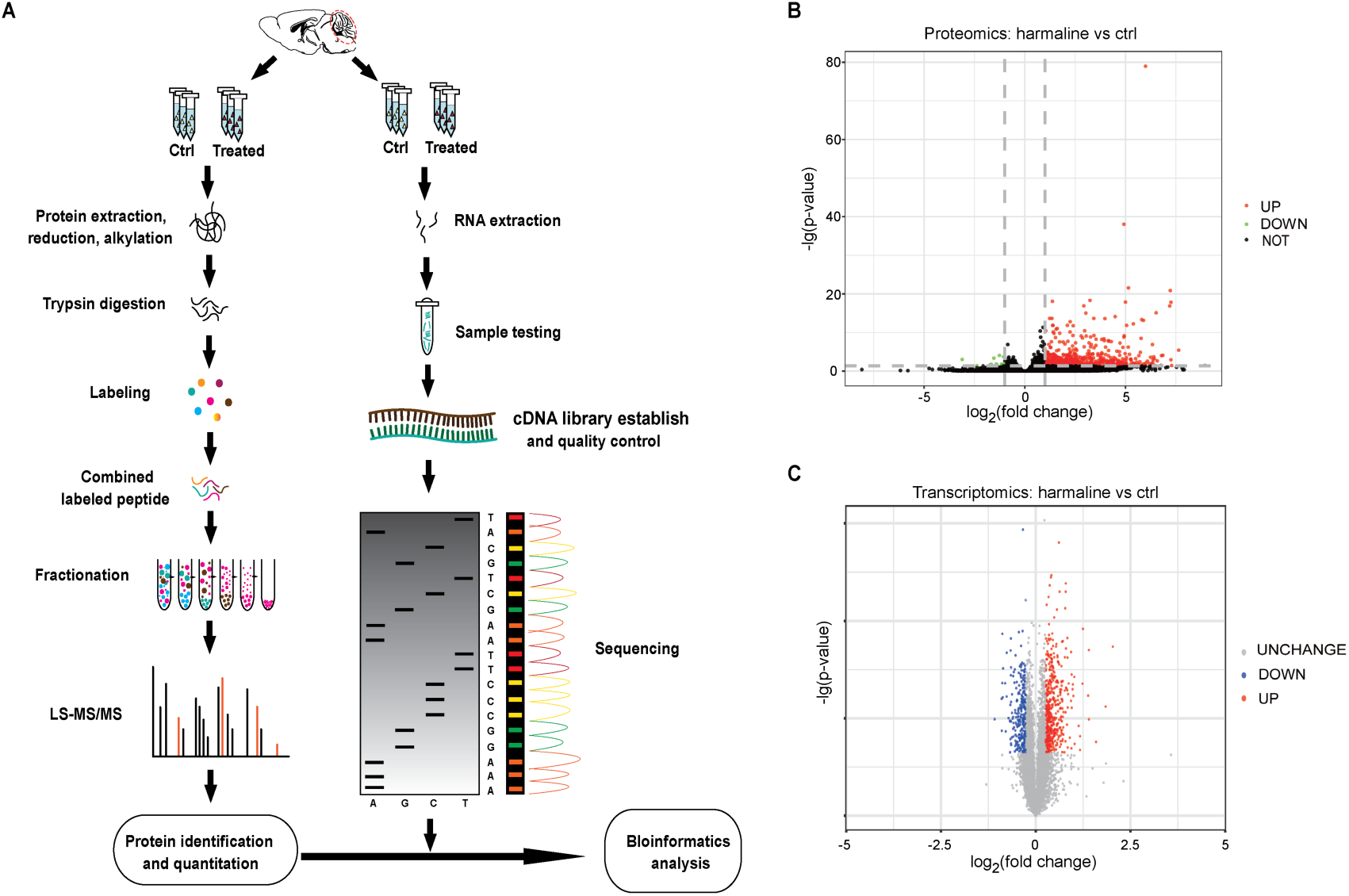
Transcriptomic and proteomic workflow and preliminary screening of candidate genes involved in harmaline-ET. (**A**) Schematic workflow of RNA-seq and LC-MS/MS and joint bioinformatic analysis based on mouse cerebellar cortex. After vehicle or harmaline treatment, mouse cerebellar cortexes were dissected, followed by independent mass spectrometry and bulk RNA-Seq and combined analysis. (**B**) 5661 proteins were identified from proteomic analysis, including 469 up-regulated proteins and 271 down-regulated proteins. (**C**) 35125 genes were identified in transcriptomic analysis, including 614 up-regulated genes and 18 down-regulated genes.

In total, we identified 35125 genes from RNA-seq and 5661 proteins from LC-MS/MS. Through gene and protein difference analysis (Fig. 2B-C), it is noteworthy that in the transcriptomic analysis, 614 genes were up-regulated, 18 genes were down-regulated; in comparison, 469 proteins were up-regulated,

271 proteins were down-regulated for proteomic analysis. However, the correlation of the gene expression level between these up-regulated and down-regulated genes and proteins in harmaline-treated mice remain unclear at this point.

### Co-occurring alterations in the cerebellum of harmaline-induced ET identified through integrated analysis of transcriptomics and proteomics

We integrated transcriptomic and proteomic information derived from the same treatment, considering genes that exhibited co-directional changes in transcription and translation as the correlated ones. In this context, 5194 correlated genes were identified, among which 19 were deemed significant, all showing upregulation (Fig.3A). Subsequently, a clustering analysis of these significant correlated genes revealed a consistent pattern of upregulation across all 19 (Fig. 3B).

**Fig 3.**
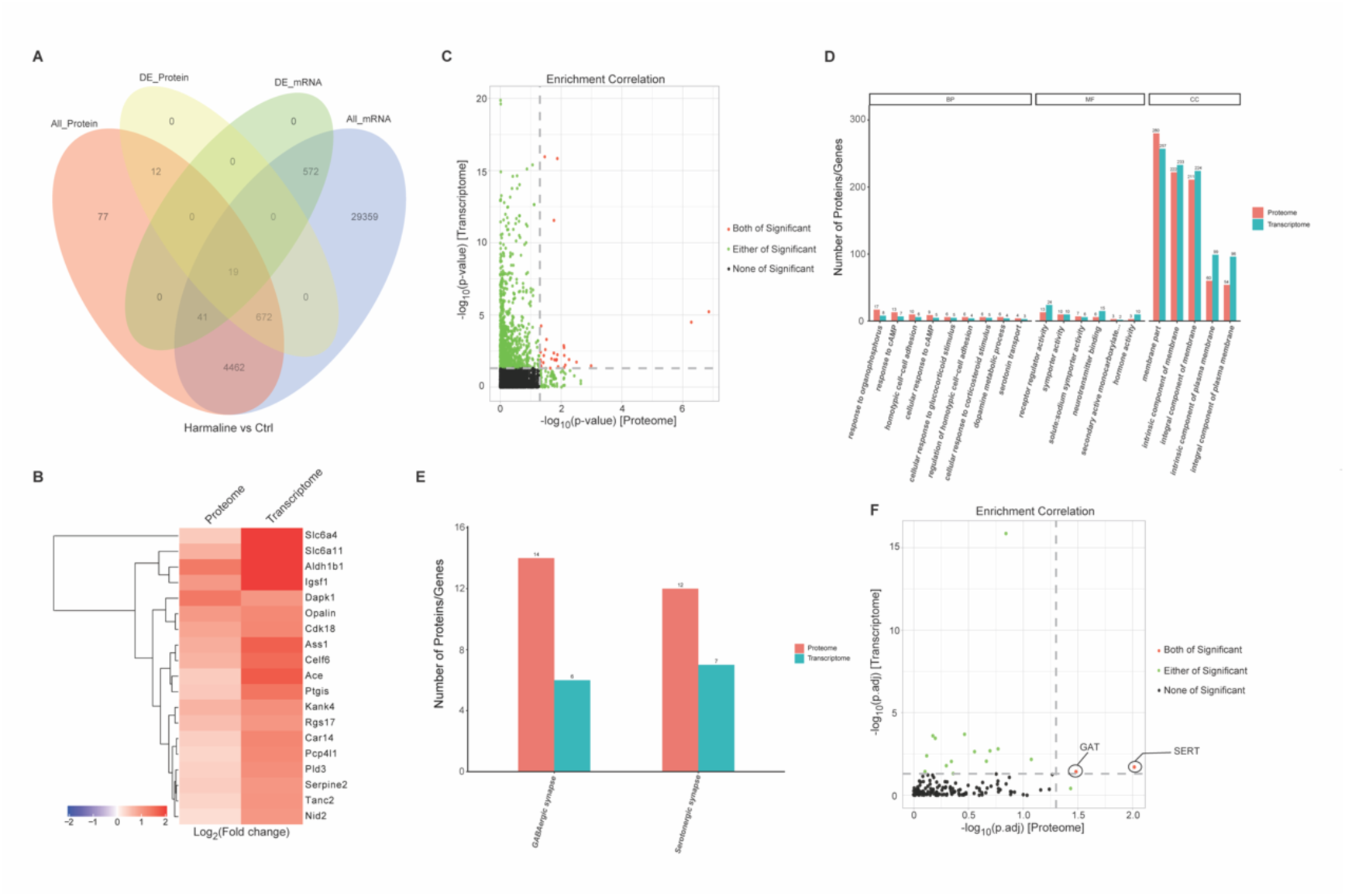
Integrated informatic analysis based on transcriptomic and proteomic data. (**A**) Integrated analysis revealed 5194 correlated genes with 19 significant upregulation. (**B**) The profile and expression richness of 19 genes based on clustering analysis. (**C**) GO analysis of enrichment correlation between proteome and transcriptome. (**D**) Combined GO analysis of enrichment correlation between proteome and transcriptome. (**E**) Combined KEGG analysis of enrichment correlation between proteome and transcriptome. (**F**) KEGG analysis of enrichment correlation between proteome and transcriptome.

To gain insights into the functional implications of those correlated genes, we conducted enrichment analysis. GO enrichment analysis identified over 30 significant GO terms that exhibiting concordance in both omics data (Fig. 3C). Notably, among these terms, proteins and genes located at the cell membrane are considered to undergo most significant changes (Fig. 3D). In KEGG enrichment, only the serotonin synapse and GABA synapse pathways stood out as highly significant in both transcriptomic and proteomic analyses (Fig. 3E). Specifically, two genes: serotonin transporter (SERT) and GABA transporter (GAT) were identified as the most significant ones (Fig. 3F). This suggests that SERT and GAT may play a significant role in harmaline-induced tremor.

### The pivotal role of serotonin transporter of cerebellar cortex in harmaline-induced tremor

To verify the potential function of SERT in harmaline-induced tremor, we conducted Western Blot analysis. Using the stable mouse model of harmaline-induced ET, we observed an increase in SERT protein levels in the mouse cerebellar cortex compared to the vehicle control group, which aligns with our integrated analysis of transcriptomics and proteomics (Fig. 4A-B). However, despite an upward trend, the quantification of GABA transporter (GAT) showed no significant difference between the two groups (Fig. 4A-C), indicating that SERT may play a more crucial role in harmaline-induced tremor.

**Fig 4.**
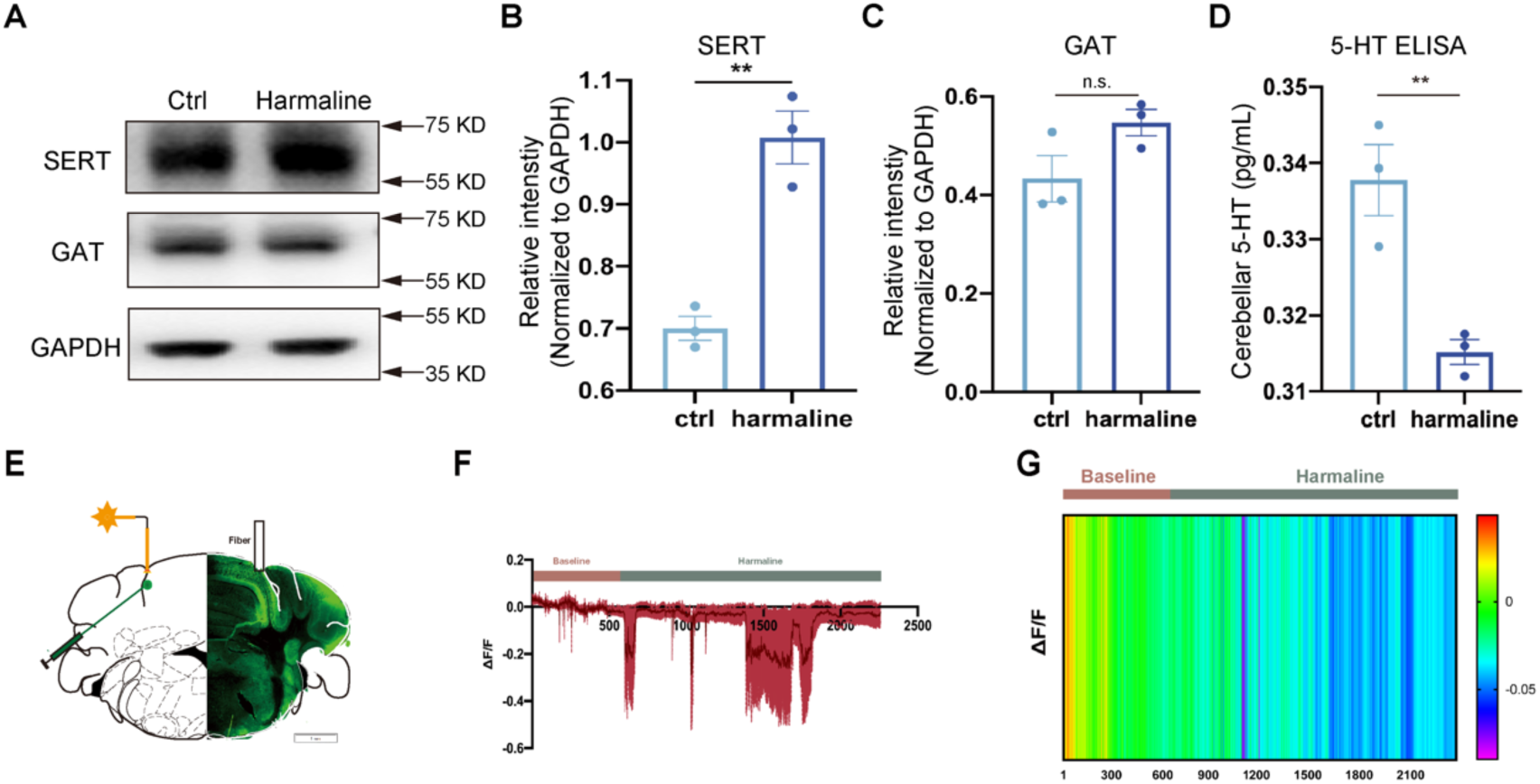
Regulation of SERT expression and 5-HT content in harmaline-induced tremor. (**A**-**C**) Representative blots (**A**) and quantification of SERT (**B**) and GAT (**C**) from cerebellar cortex in mice following vehicle or harmaline treatment as indicated. GAPDH was used as a cytoplasmic protein control. The protein level presented as mean ± SEM. For GAT, *P* = 0.1039, no significance between two groups (ctrl, 0.4331±0.0473; harmaline, 0.5471±0.268). For SERT, **P < 0.01 (ctrl, 0.7003±0.0192; harmaline, 1.008±0.426). 3 independent experiments. (**D**) ELISA quantification of 5-HT content in mouse cerebellar cortex 30 minutes after vehicle/harmaline treatment. Based on the standard curve, the 5-HT concentration: ctrl, 0.3378±0.0047; harmaline, 0.3152±0.0016. \*\**P* < 0.01. 3 independent experiments. (**E**) Schematic diagram for 5-HT sensor transfection and fiber photometry. (**F**) Quantification analysis of 5-HT sensor fluorescent fluctuation following harmaline treatment. n = 4 mice. (**G**) Representative heatmap of time-lapse recording of 5-HT sensor fluorescence.

To further investigate the relationship between SERT and serotonin (5-HT), we utilized Enzyme-Linked Immunosorbent Assay (ELISA) and found a significant down-regulation of 5-HT in the cerebellum during harmaline-induced tremor (Fig. 4D). We then locally expressed genetic sensor of 5-HT in the cerebellar cortex by AAV to continuously monitor the fluctuation of 5-HT with high temporal and spatial resolution (Fig. 4E). The results revealed that within 30 minutes of harmaline administration (intraperitoneal), the content of 5-HT declined with fluctuations (Fig. 4E, G), confirming the ELISA findings. This suggests that the down-regulation of 5-HT may be one of the consequences of the increase in SERT.

### Targeted inhibition of SERT reverses the excitability and tremor induced by harmaline

To functionally elucidate the role of SERT in harmaline-induced tremor, we conducted behavioral experiments in mice by inhibiting SERT. The mice were pre-injected with DSP-1053, an inhibitor of SERT, one hour before harmaline administration (Fig. 5A). Remarkably, with DSP-1053 pre-treatment, the harmaline-induced tremors were significantly attenuated in mice (Fig. 5B-D). These results emphasize the critical role of SERT in harmaline-induced tremor. To further investigate the potential mechanism of harmaline-induced tremor and targeted inhibition of SERT, we conducted electrophysiology experiments on cultured primary cerebellar cortical neurons. Using the whole-cell patch clamp technique, we observed that harmaline perfusion enhanced the activity of Purkinje cells (PCs) by increasing spike firing, both in amplitude and frequency (Fig. 5H-L), confirming the impact of harmaline on PC activity. Importantly, pre-perfusion with DSP-1053 significantly reversed the effect of harmaline, highlighting the involvement of SERT in harmaline-enhanced cerebellar activity (Fig. 5H-L). These results demonstrated that SERT plays a pivotal role in harmaline-induced tremor by affecting PC excitability.

**Fig 5.**
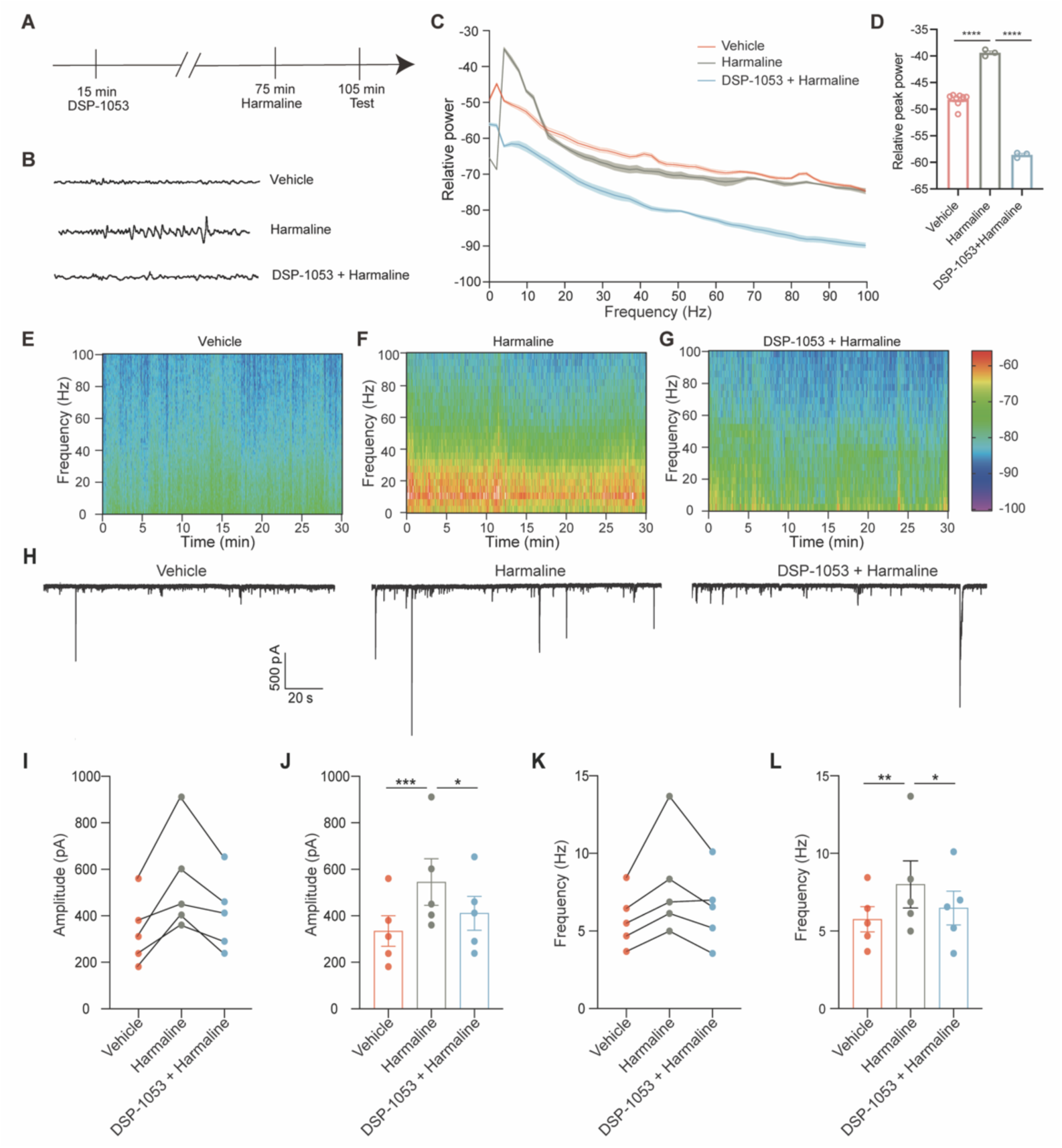
The cellular and molecular mechanisms underlying harmaline-induce tremor. (**A**) Schematic paradigm for behavioral experiments targeting SERT inhibition. (**B**) Representative raw data of head vibration using tremor detector in vehicle, harmaline with or without DSP-1053 treatment group. (**C**-**D**) Averaged power spectrum from data in (**B**). n = 8 for control group, n = 3 for harmaline-exposure group and n =3 for harmaline with DSP-1053 group. (**D**) Comparison of averaged peak power from 8 to 20 Hz among control (n = 8), harmaline (n = 3) and harmaline with DSP-1053 group. Data presented as mean ± SEM, Ctrl, -48 ± 0.4252; Harmaline, -39.82 ± 0.6153; Harmaline+DSP- 1053, -58.6±0.2877. \*\*\*\**P* < 0.0001, one-way ANOVA. (**E**-**G**) Representative spectrograph calculated from 30-minute tremor measurements based on head vibration respectively in control group, harmaline group and harmaline+DSP- 1053 group. (**H**) Representative traces of spontaneous postsynaptic current (sPSC) on cultured primary cerebellar neurons at DIV10. (**I**-**J**) Amplitude quantification of sPSC. (**K**-**L**) Frequency quantification of sPSC. n = 4 cells. \**P* < 0.05, \*\**P* < 0.01, \*\*\**P* < 0.001, one-way ANOVA.

## Discussion

The mounting evidence implicates alterations in cerebellum in ET. Postmortem examinations of ET patients have revealed characteristic neuropathological changes, including torpedo-shaped swelling of neuronal branches^20^ and decreased density in Purkinje cells^21^, as well as changes in hairy baskets for basket cells^22^, among others. In laboratory animals, harmaline-induced tremor serves as a recognized model for studying ET, providing a valuable tool for qualitatively assessing the effectiveness of pharmaceutical interventions. However, the potential mechanisms underlying harmaline-induced tremor remain largely unknown.

Our approach, which combines proteomic and transcriptomic sequencing technologies, introduces a novel dimension to elucidating gene signatures by reducing the false discovery rates inherent in individual technologies and utilizing a more precise bioinformatics infrastructure^23,24^. Within our dataset, we observed significant up-regulation of 19 genes along with their associated proteins. Through KEGG pathway analysis, we identified two specific genes, GAT and SERT. However, our Western blot results revealed a disparity between the two. Unlike SERT, the quantity of GAT did not show a significant increase after harmaline treatment. This discrepancy may be attributed to the lower precision of WB, as it only reflects proteins with marked variations at the protein level.

SERT, a component of the serotonin system, is known for its involvement in various neuropsychiatric disorders, including depression, bipolar disorder, anxiety, and neurodegenerative conditions^25^. The structural similarity between β-carboline alkaloid and serotonin has been reported^26^. However, the role of serotonin system in harmaline-induced tremor, particularly in olivocerebellar function, remains a highly debated topic^27-30^, with limited research highlighting its role in ET. Our research group made a pioneering discovery by revealing a conspicuous up-regulation of SERT in harmaline-induced tremor, suggesting its potential as a key regulatory factor in this context.

To strengthen the validity of our findings, we employed multiple experimental methods including WB, ELISA, and optical fiber recording. The collective results consistently confirmed that harmaline induces the up-regulation of SERT, leading to an increase in serotonin uptake and subsequently causing a decrease in cerebellar serotonin levels. Furthermore, we conducted electrophysiological assessments on cultured primary cerebellar cortical neurons and observed tremors in mice. These experiments demonstrated that the excitability of neurons and harmaline-induced tremors can be significantly suppressed. This further validated the critical role of SERT in harmaline-induced tremor. To provide additional support for this conclusion, we utilized DSP-1053, an inhibitor of SERT, and observed its effect on tremor suppression. The application of DSP-1053 further confirmed the involvement of SERT in harmaline-induced tremor.

Our findings align closely with previous studies that have suggested the potential effectiveness of serotonergic agonist, such as trazodone, in ameliorating ET^31-34^. While significant improvements were observed in only two small clinical trials (involving 2 patients^32^ in one study and 5 out of 6 patients^33^ in another), our results make a contribution to the increasing body of evidence supporting the involvement of the serotonin system in ET. These collective findings emphasize the potential relevance of targeting the serotonin system as a therapeutic approach for ET.

Nonetheless, some published articles have presented contrasting views. For instance, citalopram, a selective serotonin reuptake inhibitor, has been reported to augment harmaline-induced tremor^35^. Additionally, after harmaline injection, elevated level of 5-HT level was observed in striatum, cortex, hypothalamus, hippocampus ^36,37^ and brainstem^35^. Furthermore, sertraline escitalopram, one of the SSRI, was implicated in inducing movement disorders, including dystonia, akathisia, parkinsonian symptoms in a 38-year-old male patient^38^.

Several factors may contribute to the discrepancies noted above. Firstly, previous studies did not specifically focus on serotonin level in the cerebellar cortex, where our findings revealed a decline in 5-HT. Moreover, the observed augmentation in climbing fiber reuptake of 5-HT matches with the elevation of 5-HT in brainstem. Additionally, citalopram exihibits low affinity for various receptors^35^, including SERT, dopamine receptors and monoamine oxidase inhibitor receptors. As the citalopram dosage rise, the inhibition of these receptors could contribute to the augmentation of tremors. Conversely, it is theorized that SSRI may lead to a further reduction of serotonergic activity in brainstem, enhancing glucose metabolism and adenosine, eventually bringing about tremors^35^.

### Conclusion and perspective

The current study employed transcriptomic and proteomic analysis to elucidate alterations within cerebellar cortex following harmaline treatment in mice. Our findings confirm the up-regulation of SERT, a component of the serotonin system, in harmaline-induced tremor. This highlights a potential role for SERT in the pathogenesis of ET, offering a novel research strategy and identifying a potential molecular target for further investigation. Moving forward, several avenues for further research are suggested. Firstly, it is crucial to demonstrate the molecular pathway of SERT in the pathophysiology of ET. Secondly, exploring the potential of 5-HT in peripheral blood as an auxiliary indicator for the diagnosis of ET could enhance diagnostic approach. In addition, considering the widespread use of 5-HT reuptake inhibitors in depression, a similar therapeutic targeting SERT could be investigated for its applicability in ET treatment.

